# Nairobi Sheep Disease Virus: a historical and epidemiological perspective

**DOI:** 10.1101/2020.04.13.040014

**Authors:** Stephanie Krasteva, Manuel Jara, Alba Frias-De-Diego, Gustavo Machado

**Author notes:** **Correspondence** Dr. Gustavo Machado.

## Abstract

Nairobi Sheep Disease virus (NSDv) is a zoonotic and tick-borne disease that can cause over 90% mortality in small ruminants. NSDv has historically circulated in East Africa and has recently emerged in the Asian continent. Despite efforts to control the disease, some regions, mostly in warmer climates, persistently report disease outbreaks. Consequently, it is necessary to understand how environmental tolerances and factors that influence transmission may shed light on its possible emergence in other regions. In this study, we quantified the available literature of NSDv from which occurrence data was extracted. In total, 308 locations from Uganda, Kenya, Tanzania, Somalia, India, Sri Lanka and China were coupled with landscape conditions to reconstruct the ecological conditions for NSDv circulation and identify areas of potential disease transmission risk. Our results identified areas suitable for NSDv in Ethiopia, Malawi, Zimbabwe, Southeastern China, Taiwan, and Vietnam. Unsuitable areas included Democratic Republic of Congo, Zambia, and Southern Somalia. In summary, soil moisture, livestock density, and precipitation predispose certain areas to NSDv circulation. It is critical to investigate the epidemiology of NSDv in order to promote better allocation of resources to control its spread in regions that are more at risk. This will help reduce disease impact worldwide as climate change will favor emergence of such vector-borne diseases in areas with dense small ruminant populations.

## INTRODUCTION

### Background

Small ruminant populations have become one of the pillars of socio-economic wellbeing for developing countries due to their direct contribution to food security, however newly emerging diseases pose a constant threat (1–3). The demand for small ruminant products is growing globally, especially in China, the biggest producer and importer of ovine meat (4), and now as an alternative to pork consumption due to the circulation of African Swine Fever (5). Other countries such as the United States, Islamic Republic of Iran, Japan, and Qatar have also increased import demand for these products (4). Some countries such as Somaliland have economies that depend entirely on their livestock industries and employ over 70% of the population (6,7). Others such as Saudi Arabia import 5 million live ruminants a year, most of which are sheep and goats (8). However, this economic value is directly dependent on animal health, which has been compromised in regions with high incidence of tick-borne diseases (9–11). It has been proposed that the effects of ticks and tick-borne diseases on livestock pose the greatest barrier to economic development (12–14), and thus, a better understanding of current emerging disease distributions is important for human socio-economic development and animal welfare.

Some of the most pathogenic diseases of small ruminants include viruses from the *Bunyaviridae* family, which are spreading to novel areas and currently threaten small ruminant populations worldwide (15–18). There are seven serogroups within the *Nairovirus* genus, with the most impactful ones being i) Crimean-Congo hemorrhagic fever group which includes the human pathogen Crimean-Congo hemorrhagic fever virus (CCHFv), and ii) Nairobi Sheep Disease group that contains Nairobi Sheep Disease virus (NSDv) and Dugbe virus (19). Due to the similarities between NSDv and CCHFv, studies on NSDv will be useful in furthering our understanding of this important human pathogen (20), and will serve as a good model system to study other nairoviruses (21).

NSDv is characterized by hemorrhagic gastroenteritis, fever, abortion, and high mortality in small ruminants (22) and febrile illness, nausea, vomiting, and headache in humans (23). Mortality rates in susceptible animals exceed 90%, causing significant economic losses for production systems (22). This disease is listed as notifiable to the World Organization for Animal Health (OIE) (24), and it has the potential to impose trade barriers and consequently have a substantial impact on small ruminant producers worldwide (25).

Although the occurrence of NSDv has been reported in numerous countries (16,26–32), there has been limited understanding of the biogeographic factors shaping its distribution and the potential areas at risk for future epidemics. The available literature on NSDv ranges from 1910 to 2019, with host serology and virus isolation found in 14 of these studies (Table S1 in Supplementary Material). It has been found in environmentally varying areas, leaving an open question regarding the requirements of this virus and its vectors to efficiently spread infection. For that reason, the aim was to develop a systematic review and distribution model of NSDv.

### NSDv vectors and affected species

Ticks are known to transmit a greater variety of pathogenic microorganisms than any other arthropod vectors, and are among the most important vectors affecting livestock and humans (33–37). NSDv spreads via feeding of competent infected ticks, and its geographic distribution is therefore limited to the areas comprising suitable environment conditions for them (20). The main tick species related to the spread of NSDv are *Rhipicephalus appendiculatus* in East Africa and *Haemaphysalis intermedia* in Asia (20). Lewis (38) demonstrated that *R. appendiculatus* can potentially retain NSDv for long periods (138 to 871 days) depending on the life stage of the tick. NSDv has also been isolated from *Amblyomma variegatum* (39), *Rhipicephalus hemaphysaloides* (40), and *Haemaphysalis longicornis* (16).

NSDv is also known to have the potential to infect humans. The first human infection was recorded in a young boy (41) and some laboratory acquired infections occurred in following years (42,43). Human sera has been shown to contain antibodies against NSDv in India (41,44), Uganda (29), Kenya (45), and Sri Lanka (32). Other isolates have also been obtained from different vector species such as *Culex vishnui* mosquitoes, where the virus was unable to replicate without a tick host (46), *Haemaphysalis wellingtoni* ticks feeding on red spurfowl (*Galloperdix spadicea*) (47), and from a pool of *Culicoides 23* midges (48) and *Culicoides tororensis,* although this could be due to blood meal residues in the midges (49). Low titers were found in various wild ruminants, but this was likely due to an antibody cross-reaction with other viruses (28).

### The emerging pathway of NSDv

The first reports of NSDv came from Nairobi livestock markets in Kenya in 1910 as the result of an investigation carried out by veterinary pathologist Eustace Montgomery (27). NSDv was first identified as the causative agent in a classic study that showed virus transmission transovarially and transstadially in the Ixodid tick *R. appendiculatus* (27), findings that were later confirmed by Daubney and Hudson (50). In the following years, it was identified in various other areas of Kenya (28,39,51), Uganda (22,29), Somalia (attributed to *Rhipicephalus pulchellus*) (30), and Tanzania (52). NSDv is also known as Ganjam virus in Asia. Similarities between NSDv and Ganjam virus were highly debated due to their occurrence on different continents and association with different vectors, but genetic and serologic analyses demonstrated that Ganjam virus is an Asian variant of NSDv, instead of a different virus as previously described (23,52–54). NSDv was first isolated on the Asian continent in 1954 from *H. intermedia* in India (26) and Sri Lanka in 1996 (32). More recently, Gong et al. (16) were the first to discover NSDv present in *H. longicornis* ticks in China. Small ruminants bred in endemic areas do not appear to be affected by the virus (20), which could be due to the presence of maternal antibodies that provide sufficient protection until the animal’s own immunity can be established (20).

In August 2017, *H. longicornis* was found in the United States for the first time in all three of its life stages, infecting an Icelandic sheep in Hunterdon County (New Jersey) that had not been transported or had any contact with foreign animals (55). The following year, this tick was found in seven other states along the Eastern US and Arkansas (56), and its presence was verified by reexamination of archived historical samples, confirming that *H. longicornis* was present in West Virginia in 2010 and New Jersey in 2013 (56). Over the past 30 years, the US Department of Agriculture has identified this tick at least six times from imported horses in quarantine (57). *H. longicornis* is unique for its parthenogenesis, which allows a single female to produce a clonal population without mating with a male (58). *H. longicornis* is also found in Australia, which may facilitate the appearance of NSDv in the country (59). Due to the potential global distribution of this disease, the Australian Veterinary Association has stated that changes in climate could lead to an increase in disease incidence with devastating consequences (59). *H. longicornis* is the only exotic tick established in New Zealand that has an economic impact on livestock (60) from which NSDv was most recently isolated from in China (16). The range of NSDv and its vectors is likely spreading, and it will become even more important as we continue to push for breed improvement and maximizing land use to manage increasing global demands for small ruminant products (21).

### Variables affecting NSDv spread

Climate change is known to impact the distribution of a great number of diseases (61–65), especially in tropical and subtropical regions (66–68). Our understanding of the environmental preferences facilitating the survival of NSDv vectors and hosts will allow at-risk regions to better prepare for potential incursions of this disease. These tick species mentioned above are known to span large geographical ranges, such as *R. appendiculatus,* which has a territory that extends from the tropical regions of East Africa to the temperate regions of South Africa (69), as well as India and Pakistan (70), or *H. longicornis*, which occupies a wide range of climates from equatorial New Guinea and the Pacific Islands, to snow and cool summer conditions in northeast Primorsky Krai, Russia (71,72). For that reason, a deeper understanding of the environmental conditions facilitating the survival of these ticks is key to predicting novel areas at risk for NSDv spread, which could potentially lessen the impact of this disease on naive populations.

### Economic impact of NSDv

As global trade continues to increase, infectious diseases are posing a greater threat to our economies, food supply, and health (73,74). NSDv has the potential to have devastating effects on naive sheep and goat populations (22), and therefore, efforts to increase awareness and surveillance should become a priority, especially for countries with economies that rely heavily on small ruminant products. NSDv vectors such as *H. longicornis* have already adapted to New Zealand and Australia (59,60), threatening the NSD-free status of these countries. Crossbreeding and the introduction of new breeds of small ruminants have been the target of recent efforts to increase the efficiency of meat production and commerce in some countries. This increases the chances of NSDv outbreaks in populations lacking immunity against the virus (75–77) since the main route for NSDv spread occurs upon movement of susceptible animals into enzootic areas (51). Since small ruminants are vital to people’s livelihoods in low-income communities (1), efforts to preserve their health and better understand tick-borne diseases are essential.

Given the critical importance of small ruminant production to global food security and the potential for NSDv spread in new regions, the current information gaps in the epidemiology of this disease have to be explored. This allows us to better predict future outbreaks and potential regions at risk for this disease. This is the first systematic review and predictive model of NSDv, allowing us to assess the potential distribution of the disease and predict regions that are most at risk.

## METHODS

We divided this study into two main sections: i) a systematic review to identify all susceptible hosts, potential vectors, and the ecological dependencies for virus spread and ii) an assessment of the geographic and environmental distribution of hosts and vectors using an ecological niche modeling approach.

### Systematic review

#### Literature selection

We followed the steps outlined by O’Connor and Sargeant and others (78–80) (Table S2 in Supplementary Material) to scope the global distribution of NSDv. Our search terms were designed to recover all information and reports under the terms “Nairobi Sheep Disease” and “Ganjam virus”. The study protocol followed the Preferred Reporting Items for Systematic Reviews and Meta-Analyses (PRISMA) guidelines (81,82) (Figure 2).

**Figure 1.**
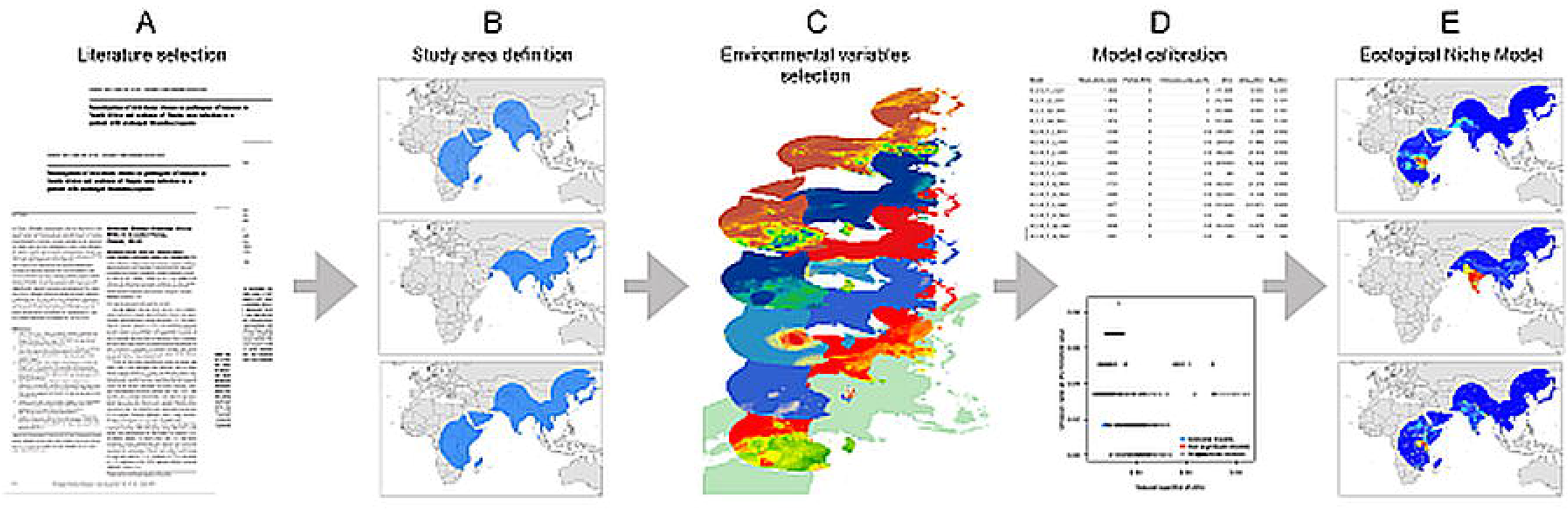
Workflow of the modeling process. **A)**Data collection based on literature review, **B)**Study area definition, **C)**Environmental variables selection, **D)**Model calibration, and **E)**Final ecological niche model.

**Figure 2.**
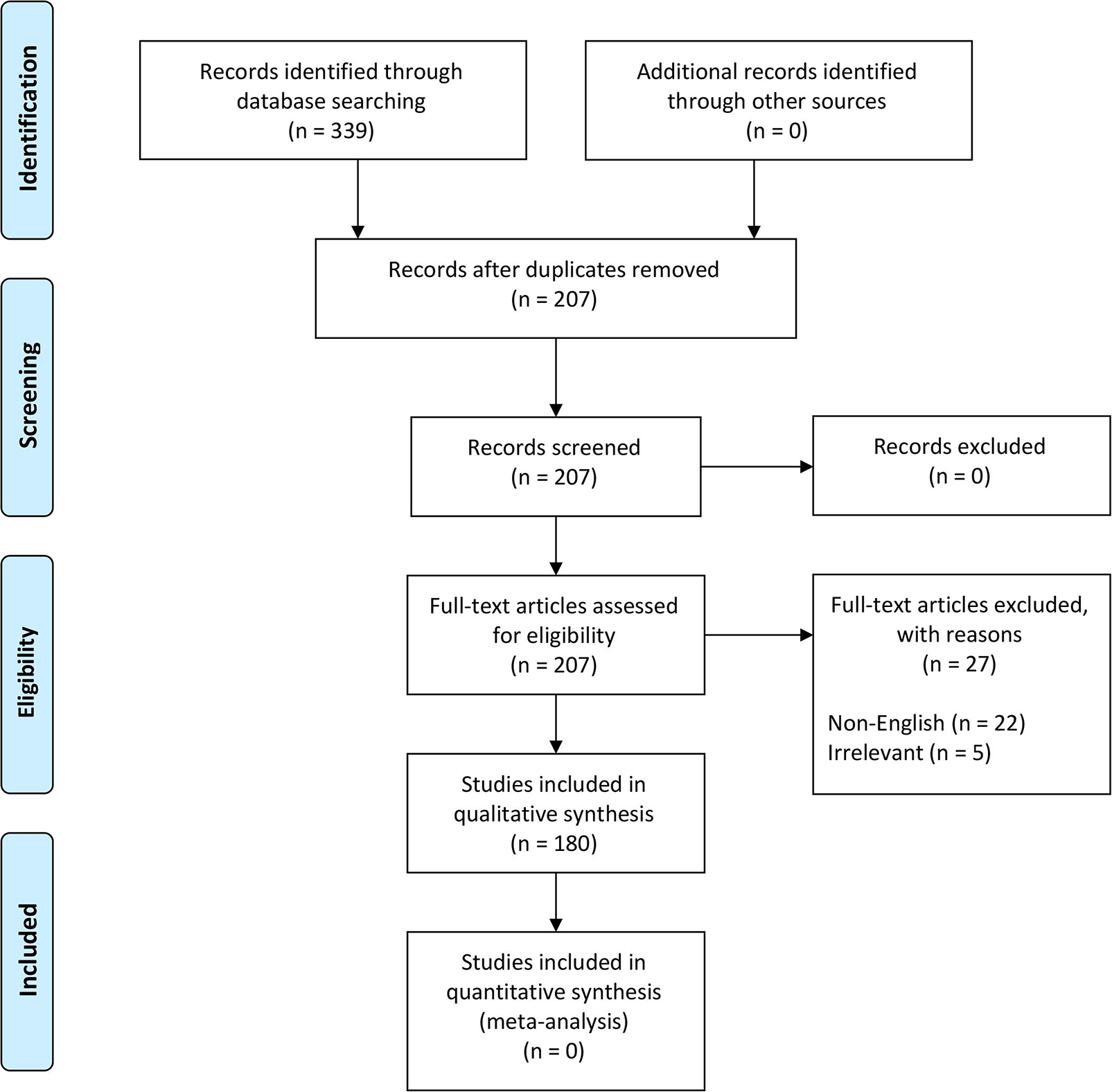
PRISMA flow diagram detailing the literature selection process (81).

These terms were used to search all databases using the EBSCOhost platform, which yielded 339 scientific publications in 11 databases (Table S2 in Supplementary Material). Exclusion criteria included duplicates (132), non-English papers (22), and irrelevant studies (5) which brought the total number of eligible papers for systematic review to 180 (Figure 2). This database search was completed in November 2018.

### Ecological niche model

#### Data collection

From the literature review, we extracted a total of 308 locations from 14 studies. The information retrieved included occurrence data classified as follows: i) host occurrence- if the virus was identified from sheep, goat, or human serology, ii) tick occurrence- if the virus was identified directly from tick vectors. Data points were georeferenced to the centroid of the smallest administrative division of each country from where the data was collected. One exception was the use of the locations reported by Gong et al. (16), which were also verified before being considered for analysis. Diagnostics based on post-mortem examinations were not considered since lesions are not specific enough to reliably distinguish NSDv from heartwater, *Pasteurella* pneumonia, or babesiosis (30). Additional occurrences from the following studies in South Africa (83), Uganda (29), Kenya (39), and India (41) were not included in the study due to non-specific locations from which we were unable to recover the centroid of the smallest administrative boundary.

#### Model calibration area

To define the model calibration area for the circulation of NSDv, we followed the framework proposed by Soberón and Peterson (84), which restricts the ecological niche model to ecological features for the organism in question, the resolution of the environmental variables employed, and the extent of the region where the organisms are able to disperse due to their biogeographic barriers (see M in the BAM framework in Soberón and Peterson (84)). This region is defined as the area accessible to the species that has been sampled, so presence records can exist within a suitable area (85). We measured the maximum distance between all NSDv locations and used this distance as the accessible area for the virus. A buffer around the occurrences was applied to establish the model calibration area.

#### Environmental variables selection

The environmental variables used to estimate the distribution of NSDv were selected based on the described requirements of the hosts and ticks, including their survival in landscapes with suitable temperature and humidity and their amplification by the presence of livestock reservoir species. We used six environmental variables to calibrate all models, and candidate models were selected based on the described requirements of the virus, including its capacity to survive in the landscape based on the presence of susceptible host species. We included the variables shown in Table 1, which will serve as an appropriate approximation of the required biogeographic conditions.

**Table 1.**
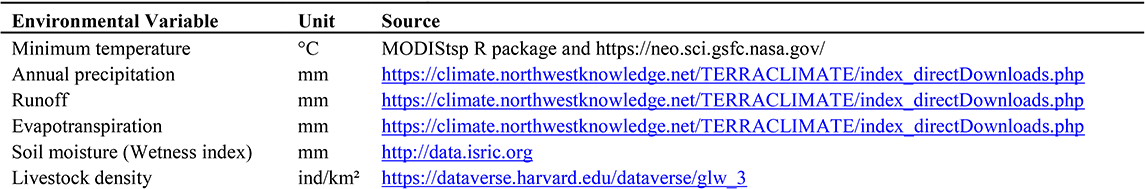
Variables used for the NSDv ecological niche models.

Previous studies have highlighted the importance of temperature for the main host of NSDv, *R. appendiculatus* (69,86,87). Although it is not a critical limiting factor for ticks, it is still important, especially when it approaches the developmental threshold for that species (71). Likewise, variables related to humidity are also important because it has been observed that prolonged drought and dry pastures cause high mortality, especially among engorged *H. longicornis* (71,86) and *R. appendiculata* larvae and eggs (87), and *H. intermedia* and *R. haemaphysaloides* numbers which seem to increase after rains (88). For that reason, we used precipitation, runoff, evapotranspiration, and soil moisture as predictors of tick prevalence. Finally, livestock population was represented by the density of sheep and goats, which are most clinically affected by NSDv (53,89).

#### Ecological niche model

Niche modeling presents a framework based on ecological theory by which to interpret, understand, and anticipate geographic distributions of species and other biological phenomena, such as disease transmission (90). For this study, we considered an ecological niche as the set of environmental conditions in a region necessary for a species to persist (85). Information about the presence of susceptible hosts for NSDv were also added to this study through a layer that contained sheep and goat densities. We explored three scenarios based on different combinations of occurrences: i) the occurrences of ticks that tested positive for NSDv; ii) the occurrences of sheep and goats that tested positive for NSDv (hosts); and iii) the combination of both positive tick and host occurrences. For these analyses, we considered that the factors explaining the distribution of NSDv and its potential hosts follow the Biotic-Abiotic-Mobility (BAM) framework (84). The analysis was set individually for each occurrence data set for all models in which a hypothesis for each accessible area M was constructed (further details of the importance of this step can be found in Barve et al. (85)). The study area for positive ticks and host cases was designed based on the union of buffer areas created around each occurrence location. The buffer was built based on the average distance between the external points and the most central point (centroid). Thus, for positive ticks this buffer was defined as 1,334 km, while for positive hosts, it was 1,555 km. The polygons drawn around each study area were then used to intercept the environmental predictor layers (Table 1).

The total occurrences for each set were randomly subdivided into 70% of the data set for model calibration and 30% for model evaluation. This data division allowed for both model calibration and internal testing. For each combination of data, we created 493 candidate models by combining 17 regularization multipliers (0.1-1.0 by the interval of 0.1, from 1-10 at the interval of 1), with all 29 possible combinations of the five MaxEnt feature classes (linear=L, quadratic=Q, product=P, threshold=T, and hinge=H). We evaluated and selected the candidate model performances based on significance rates (5%), and model complexity penalizations (AICc). The best models for each dataset were selected according to the following criteria: significant models with omission rates <=5%, and from those selected models, we used delta AICc values of <=2 to determine the final candidates. Both the full model calibration and selection step were performed using the “kuenm” package in the R environment and used Maxent as the modeling algorithm (91).

The final ENM models were performed in Maxent version 3.3.3 k (92). The specific configuration of all models included 10 bootstrap replicates and random seed with logistic outputs. Finally, to identify extrapolation risk in the model transfer steps, we used the mobility-oriented parity (MOP) index for each data set, which is an improved metric proposed by Owens et al. (93). The interpretation of the output models followed Merow et al. (94) as a suitable index to account for the probability of disease risk. To further evaluate model predictions, continuous outputs were converted into binary maps based on a threshold, removing 10% of the calibration occurrences (error = 10%) to reduce the uncertainty of estimations and facilitate model interpretation (95). Finally, the models were transferred to a global scale applying the same predictor layers used for the calibration step (Table 1).

## RESULTS

### Descriptive Analysis

A descriptive analysis of both the disease and vector distribution is provided in Figure 3, even though the total number of recovered disease occurrences were not used as offset information. A total of 308 data points were extracted from 14 studies, from which 12 were isolated from ticks and the rest were obtained from host serology. Data coming from hosts was distributed throughout East Africa, India, and Sri Lanka, while tick data was found only in India, Sri Lanka, and China.

**Figure 3.**
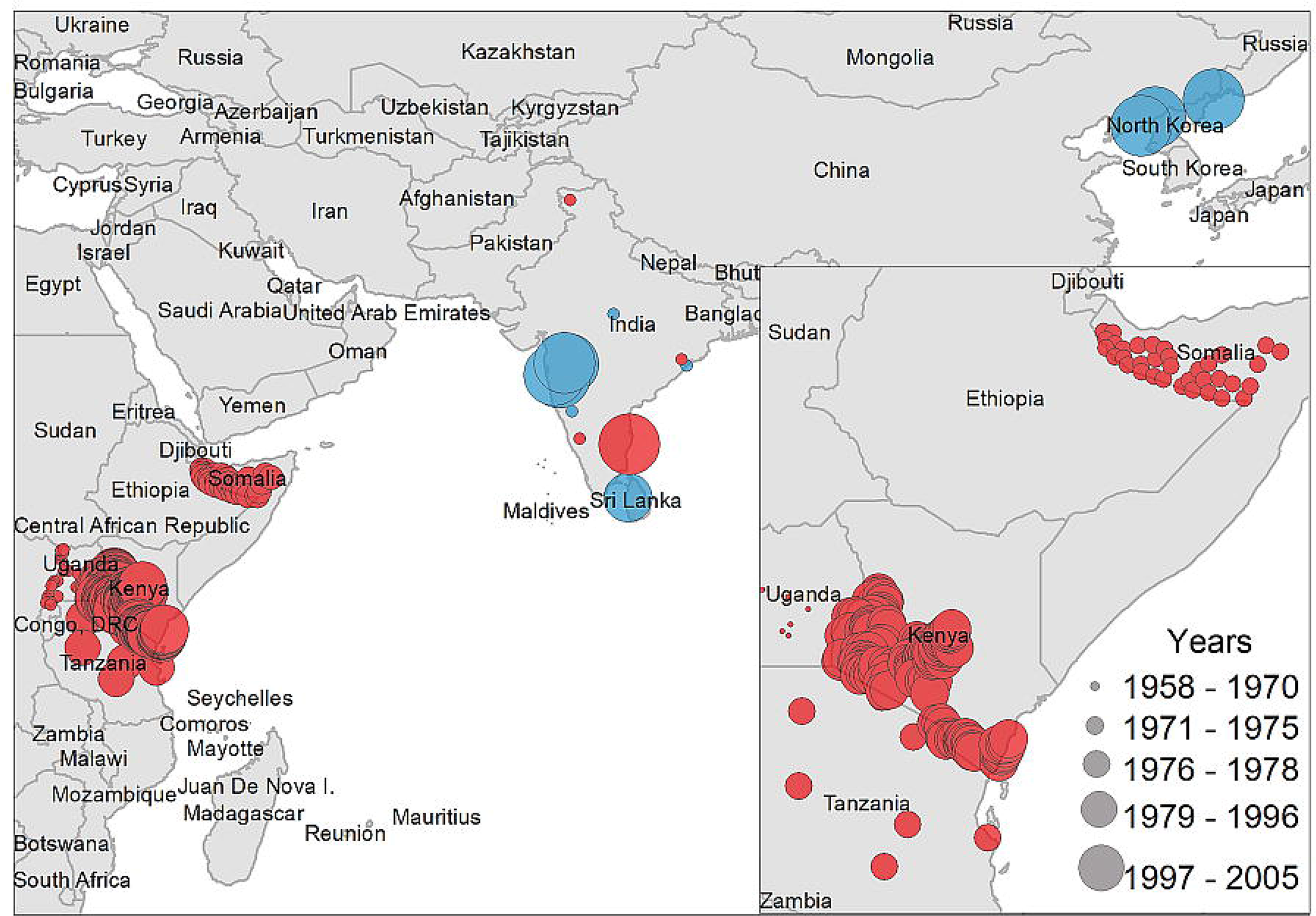
Case distribution of NSDv used for model calibration. Points denote occurrence locations. Color represents positive host (red) and tick (blue) occurrences and size denotes the year of publication.

### Ecological niche models

#### NSDv host potential distribution

The positive host occurrences of NSDv were used to calibrate 493 models along with the full set of environmental predictors. In total, 490 candidate models were significant from which 157 were significant and met the omission rate criteria. For the global minimum AICc values, 1 model had delta AICc values <=2, and met the full criteria used in the selection step (Figure 4). We emphasize that this selected model met both criteria but did not have lower AICc, which is often utilized as single decision criteria in model selection (96–98) (see the blue triangle which highlights the selected final mode- Figure 4). In order to identify areas of higher risk for NSDv occurrences, the selected model identified the geographic risk areas for small ruminants. This model showed countries presenting low suitability for NSDv spread including Democratic Republic of Congo, Zambia, and southern Somalia. There were clusters of unsuitability throughout Saudi Arabia, eastern Yemen, Oman, and western China of the study area. High-risk regions were found throughout Eritrea, Ethiopia, Kenya, Uganda, Rwanda, Burundi, Tanzania, Malawi, Zimbabwe, and the southern tip of India (Figure 7). The livestock density variable was the most influential (48.7%) in describing the risk area, followed by runoff (21.8%), evapotranspiration (11.6%), precipitation (10.2%), minimum temperature (4.7%), and soil moisture (3%).

**Figure 4.**
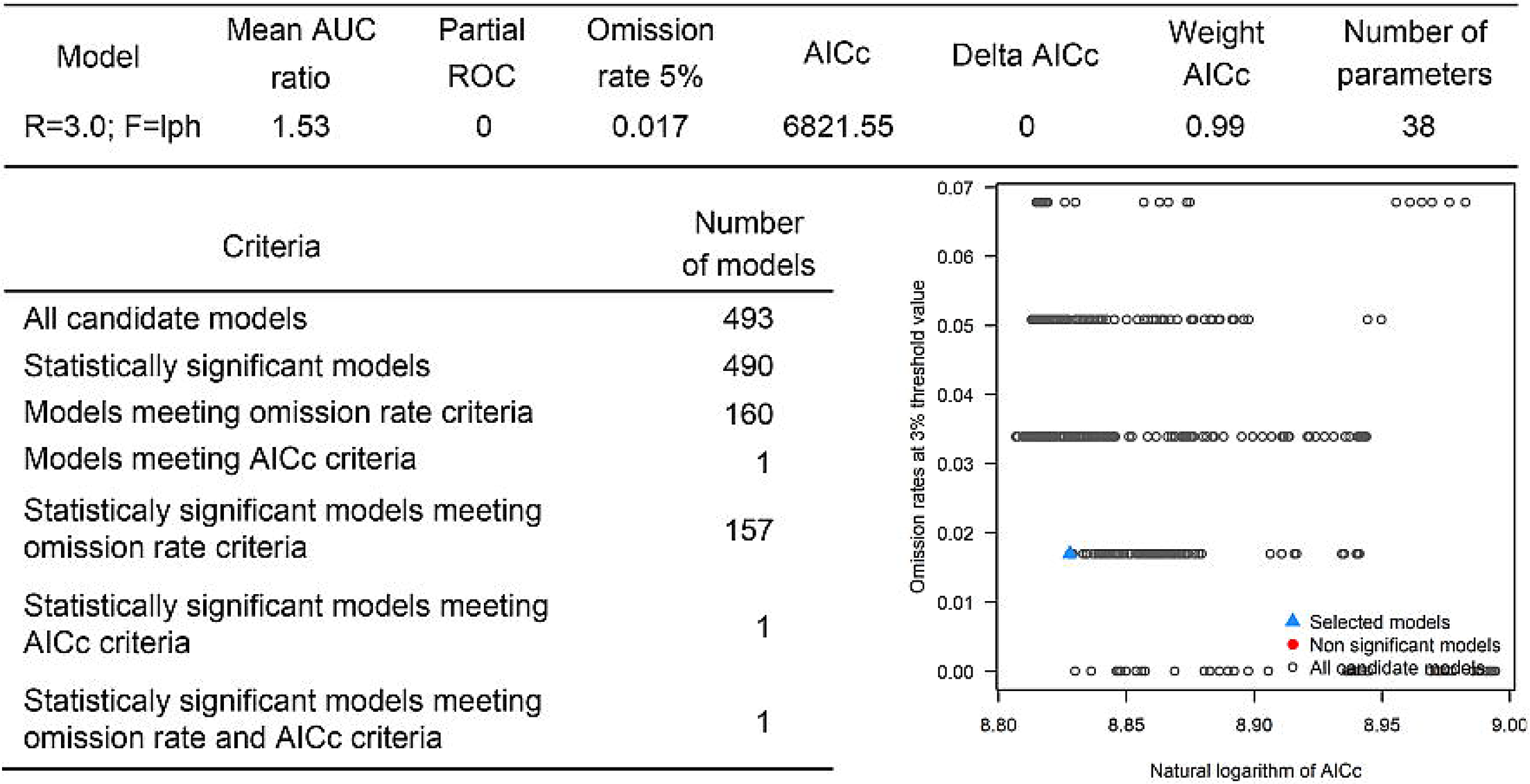
Model calibration statistics for hosts.

#### NSDv tick potential distribution

The occurrences of NSDv in vectors was modeled applying the selected parameterizations found in Table 1. Using the locations for tick occurrences, 367 candidate models out of 493 were statistically significant if compared with the assumed null expectations. All 367 of these statistically significant models also met the omission rate criteria. For the global minimum AICc values for ticks, 2 models had delta AICc values <=2 and 1 model met the criteria (Figure 5). The model selected had lower AICc, was significant, and within the omission error cutoff (see the blue triangle which highlights the selected final mode- Figure 5). This model was used to identify areas with a potential circulation of ticks within the calibration study area. It showed unsuitable areas clustered in Sri Lanka, central Vietnam, eastern Taiwan, the western shores of Japan, and southeast China. High-risk regions were found throughout the southern half of India, Sri Lanka, Bangladesh, Myanmar, Laos, Thailand, Cambodia, Taiwan, China’s Hainan Island and the southern tip of Guangdong province, and the northern and southern portions of Vietnam (Figure 7). The minimum temperature variable was the most influential (53.5%) in describing the risk areas, followed by livestock (38.4%), soil moisture (4.6%), and runoff (3.5%). Precipitation and evapotranspiration were not influential (0%).

**Figure 5.**
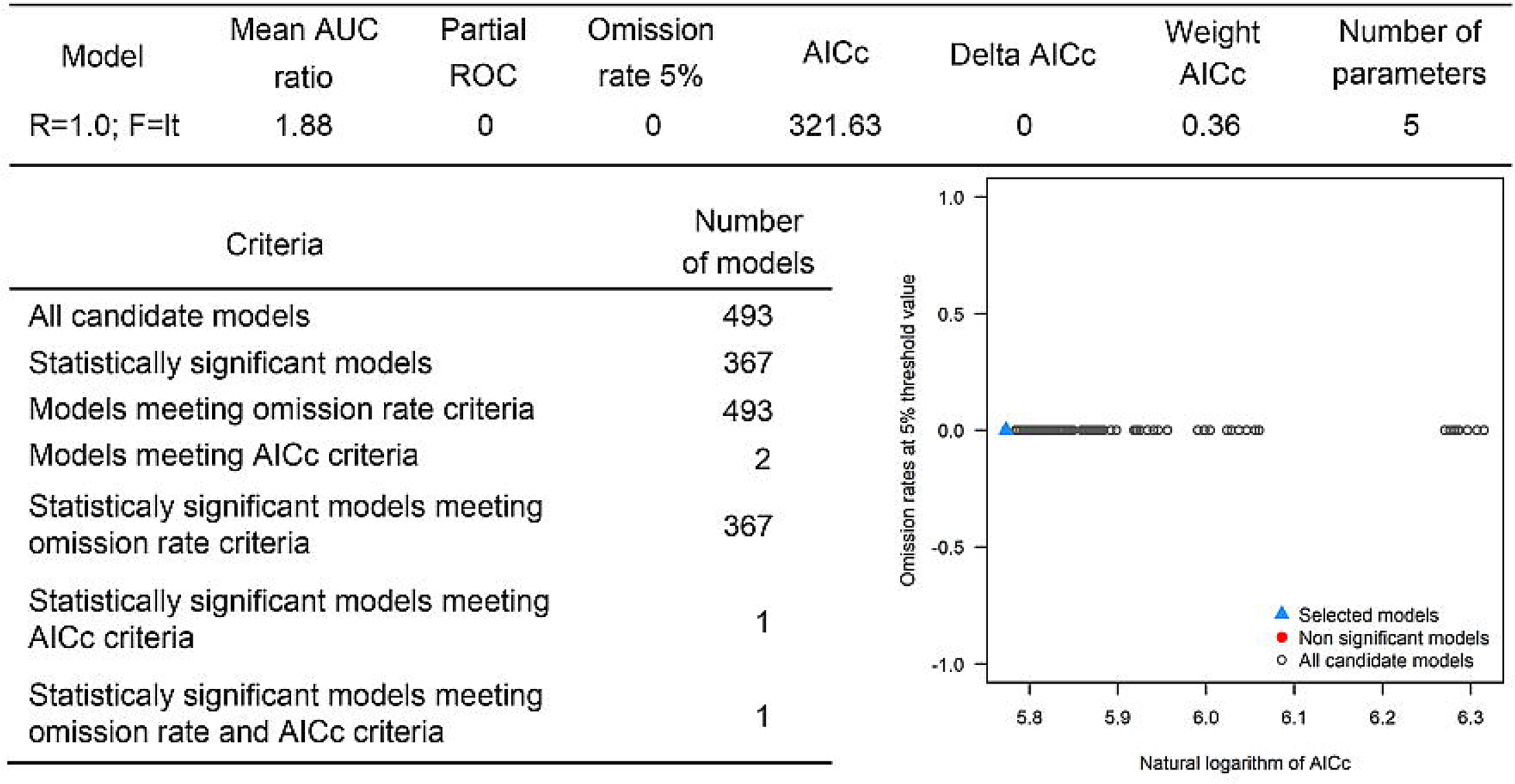
Model calibration statistics for ticks.

#### NSDv host and tick potential distribution

The last model exploration we did included the combination of data associated with infected hosts and the occurrences of potential vector ticks. From 493 candidate models, 209 were significant, from which 109 also met the omission rate criteria. For the global minimum AICc values, 1 model had delta AICc values <=2. Only 1 model met the full criteria used in the model selection step (Figure 6) (see the blue triangle which highlights the selected final model-Figure 6). Suitable areas in the study area included Ethiopia, Uganda, Kenya, Tanzania, the southern tip of India, the southeastern coast of China, and Taiwan (Figure 7). The livestock density variable was the most influential (38.1%) in describing the risk area, followed by evapotranspiration (27.2%), runoff (14.5%), minimum temperature (9.2%), soil moisture (7.8%), and precipitation (3.3%).

**Figure 6.**
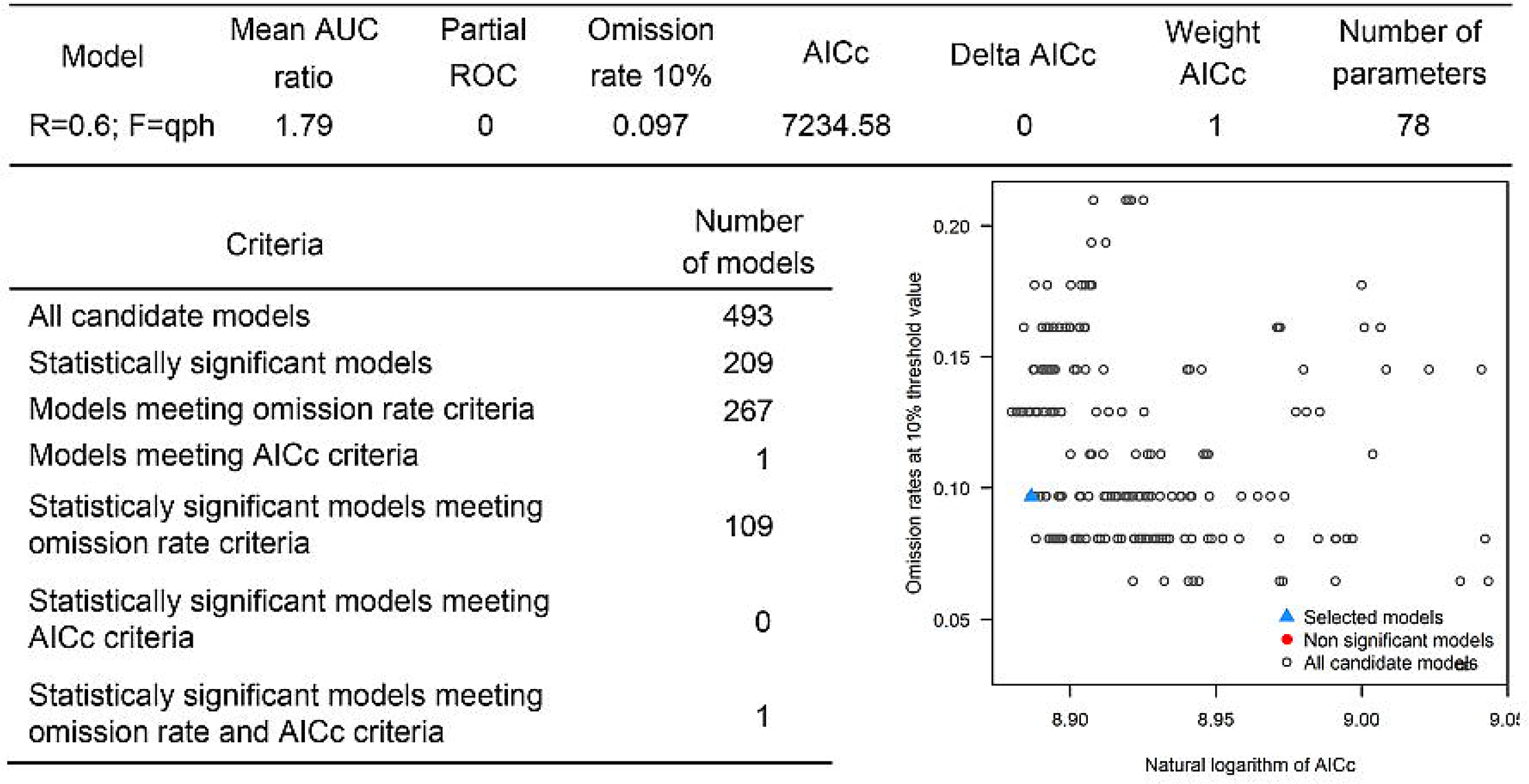
Model calibration statistics for hosts and ticks.

**Figure 7.**
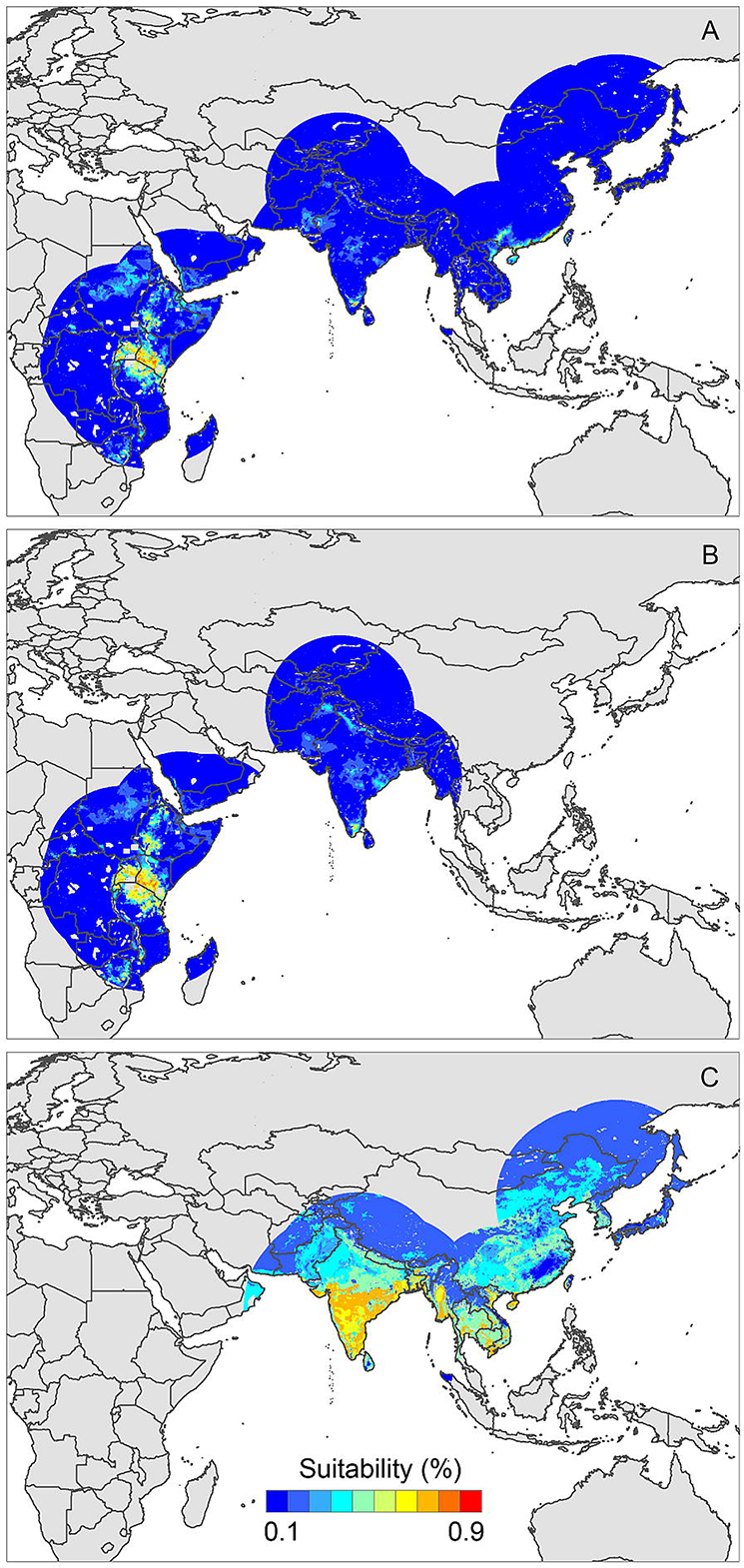
Ecological niche models of **a)**hosts and ticks, **b)**hosts and **c)**tick circulation. Warmer colors represent areas with higher environmental suitability.

#### NSDv environmental preferences

Overall, our ecological niche models coincided well with the reported geographical locations of the host and tick occurrences identified during the systematic review. These regions are the most environmentally suited for NSDv, and they indicate areas where the disease would most easily become established and continue spreading to naive small ruminant populations. For the host-based model, the most suitable areas occurred with the highest values of evaporation (46.1 mm; IQR 83.0 mm), precipitation (40.3 mm; IQR 63.0 mm), and runoff (2.8 mm; IQR 3.0 mm). In the case of the tick-based model, the most suitable areas presented the highest values of soil moisture (62.2 mm; IQR 588.0 mm) and livestock density (11812.0 lnd/km²; IQR 8953.8 lnd/km²). Finally, the model combining hosts and ticks presented the highest values of minimum temperature (12.86 °C; IQR 5.50 °C) (see Table 2 for detailed information).

**Table 2.** Environmental preference of hosts, ticks, and hosts & ticks based on their potential distributions.

|  | Statistic | Evapo. (mm) | Livestock (Ind/km<sup>2</sup>) | Min. temp. (°C) | Precip. (mm) | Runoff (mm) | Soil moist. (mm) |
| --- | --- | --- | --- | --- | --- | --- | --- |
| **Hosts** | Mean | 46.1 | 11471.6 | 12.8 | 40.3 | 2.8 | 39.6 |
|  | Minimum | 0.0 | 472.6 | -6.7 | 0.0 | 0.0 | 0.0 |
|  | Maximum | 164.0 | 247138.0 | 22.9 | 343.0 | 223.0 | 371.0 |
|  | DesvEst | 43.1 | 12683.9 | 4.61 | 48.8 | 8.2 | 50.8 |
|  | Range | 164.0 | 246665.4 | 29.6 | 343.0 | 223.0 | 371.0 |
|  | IQR | 83.0 | 9990.2 | 5.8 | 63.0 | 3.0 | 62.0 |
| **Ticks** | Mean | 24.0 | 11812.0 | 3.9 | 13.8 | 0.8 | 62.2 |
|  | Minimum | 0.0 | 7.2 | -32.0 | 0.0 | 0.0 | 0.0 |
|  | Maximum | 145.0 | 182469.0 | 23.3 | 230.0 | 107.0 | 588.0 |
|  | DesvEst | 23.1 | 9575.1 | 12.4 | 22.3 | 2.2 | 71.7 |
|  | Range | 145.0 | 182461.8 | 55.3 | 230.0 | 107.0 | 588.0 |
|  | IQR | 38.0 | 8953.8 | 15.2 | 13.0 | 1.0 | 74.0 |
| **Hosts and**<br>**ticks** | Mean | 39.7 | 10267.0 | 12.8 | 35.4 | 2.2 | 31.8 |
|  | Minimum | 0.0 | 472.6 | -0.5 | 0.0 | 0.0 | 0.0 |
|  | Maximum | 161.0 | 148964.0 | 22.2 | 343.0 | 223.0 | 440.0 |
|  | DesvEst | 41.2 | 9545.4 | 4.1 | 42.8 | 5.4 | 43.8 |
|  | Range | 161.0 | 148491.4 | 22.7 | 343.0 | 223.0 | 440.0 |
|  | IQR | 65.0 | 9434.4 | 5.5 | 58.0 | 3.0 | 52.0 |

## DISCUSSION

We identified novel countries at risk for infection with NSDv, including Ethiopia, Malawi, Zimbabwe, Southeastern China, Taiwan, and Vietnam. These findings suggest that NSDv may have a wider range than was previously thought, and therefore, these predictions can help direct surveillance efforts to those regions where NSDv could efficiently spread if introduced in the future.

Livestock diseases can result in devastating trade bans with wide-ranging consequences (7). Specifically, NSDv has the potential to easily spread to naive populations and cause significant economic losses (22). Therefore, it is imperative for at-risk countries to be aware of these risks and take necessary precautions to protect their animals, peoples’ livelihoods, local economies, and global trade partners.

To the author’s knowledge, this is the first systematic review of NSDv that has gathered all available geographic locations where this disease has occurred. Systematic reviews are increasingly being used in animal agriculture and veterinary medicine since they offer a replicable evidence-based method to identify, evaluate, and summarize primary research (78). Based on the reports obtained, we determined the countries which are currently at risk for the introduction of this disease. We performed extensive model fitting from which we selected the best performance-based model, focusing on significance rather than AIC values as our main criteria for model selection. Using significance rather than AIC allows us to choose the single best model which outperforms the rest and will give us the most realistic prediction of occurrence. We found varying results in our models, with notable differences in the environmental preferences needed for the optimal spread of the virus when host and tick-based models were compared. Because our data points included serology from hosts and virus isolation from ticks, these models should be interpreted together rather than separately. Multiple environmental variables play a role in the transmission of NSDv, and they should all be considered holistically. We support that this combination of data would more accurately predict potential regions of outbreak since the main transmission of NSDv will likely occur when both hosts and ticks coincide together in the same environment.

ENMs have been widely applied in epidemiology to assess species introduction into novel areas and disease emergence (96,99–101), or the effect of climate on disease distribution (102–105). However, previous ENM methods rely solely on the lowest AIC to identify the best fit model, which alone may not lead us to the best option for that set of occurrences (106). Significance should be used as the first criteria when filtering through candidate models, followed by performance and simplicity (91). This approach reduces the time spent on transferring these models, and uses the mobility-oriented parity (MOP) index to prevent outcome over-interpretation and identify extrapolation risks, making it a critical tool when transferring ENMs to current and future scenarios, and resulting in a more accurate prediction of species’ ecological niche as the environment changes. These calibrations have become more popular and are being applied throughout epidemiology (91,107), but our model of NSDv will be the first time that this calibrated approach has been used to model the potential distribution of a virus.

Ticks are described as the main route of NSDv transmission and therefore, they are the most likely route of infection for naive populations given that the virus can survive for such extended periods of time inside its vector (38). International trade is a known pathway for introduction of exotic ticks into new areas, which have the potential to harbor pathogens with significant consequences for human health and agriculture (108,109), highlighting the importance of increased surveillance measures and regulatory framework over animal trade-related interactions. In the tick-based model, NSDv preferred areas shown to have elevated soil moisture and a denser population of livestock. Although the density of sheep and goats was expected to have a direct influence on the number of infected ticks found in some locations, soil moisture may have facilitated the ability of ticks to survive in the field and continue transmitting the disease to other individuals. In contrast, the model based on host occurrences had elevated mean evaporation, precipitation, and runoff. This may correlate with areas of intense feed and water resources, thus concentrating more animals in one area, which will facilitate the spread of the disease between and within the population.

Even though these two models can independently explain and predict areas where NSDv is most likely to occur, the combination and overlap of the most suitable environments for hosts and ticks will represent the areas where hosts and their vectors are more likely to proliferate and coincide, greatly increasing the possibility of disease spread. Thus, we recommend the comparison and combination of these models for future epidemiological assessments in order to get the most accurate prediction of at-risk areas.

We encountered some limitations during the development of this study. First, a number of non-English papers (n=22) were excluded in our systematic review which may have included additional data for our models. We only included data points that had a specific location and excluded those that were labeled as having been collected in general areas or districts, so it is possible that additional occurrences were not recorded. Additionally, we used a centroid to approximate the location of occurrences, which entails the possibility of some data points being farther away from the center. Multiple species of Ixodid ticks were modeled together due to their similar environmental requirements, which may overgeneralize the distribution of these ticks.

Vector-borne diseases are expected to be the most climate-sensitive subset of infectious diseases (110,111), and using environmental variables is critical to predicting their distribution. Recent outbreaks of devastating ruminant diseases such as Bluetongue and epizootic hemorrhagic disease, serve as a reminder of the consequences that climate change has on vector-borne diseases (112), and illustrates the importance of using the best modeling techniques we have at our disposal in order to more accurately predict the spread of these vector-borne diseases.

## Conclusions

The data compiled here will be useful for additional spatial and environmental modeling of NSDv and its vectors, informing governments and policy makers about their prevalence and spread in order to direct surveillance efforts. Further studies would also help the development of case studies for NSDv and other infectious diseases affecting small ruminants, as well as to understand and predict where to best implement surveillance strategies, increasing response time and decreasing economic losses. Further study of the tick species shown to carry and transmit NSDv will be imperative as the distribution of these species changes with environmental factors and increasing international trade. As the ticks’ distribution changes, so too should surveillance strategies. Additionally, many countries that could potentially be at risk for NSDv have never had a cross-sectional study done on their livestock or tick species of interest, which would hinder their ability to quickly and effectively respond to a disease outbreak. As an OIE reportable disease, NSDv can cause significant trade implications for countries and could be underreported as a result. Rising global demand for these products and recognition of the socioeconomic importance of small ruminants merits further discussion of the diseases that impact them and pose a threat to their populations in other parts of the world and the communities that depend on them.

## Author contributions

SK, MJ, GM contributed to the design of the project. SK collected and curated the data used in both the systematic review and modeling exercise. MJ and GM performed the modeling section. SK wrote the manuscript. All authors reviewed and edited the manuscript.

## Acknowledgements

We acknowledge the Department of Population Health and Pathobiology at North Carolina State University for providing startup funds for G. Machado

