## Supplementary material for "Nairobi Sheep Disease Virus: a historical and epidemiological perspective": ss information

**Running title: Nairobi Sheep Disease Virus distribution**

**Stephanie Krasteva[1], Manuel Jara[1], Alba Frias-De-Diego[1], Gustavo Machado[1*]**

[1]Department of Population Health and Pathobiology, North Carolina State University, College of Veterinary Medicine, North Carolina, United States of America.

[*] **Correspondence**
Dr. Gustavo Machado


**Table S1:** Data points extracted from systematic literature review.

| Author | Type | Tick Species | Country | Locality and State | Latitude | Longitude | Host |
|---|---|---|---|---|---|---|---|
| Gong et al., 2013 | Tick | *H. longicornis* | China | Jian, Jilin | 40.8666667 | 125.566667 | Sheep |
| | Tick | *H. longicornis* | China | Jinxing, Jilin | 42.4166667 | 130.633333 | Cattle |
| | Tick | *H. longicornis* | China | Dadong, Liaoning | 40.1166667 | 124.383333 | Sheep |
| Dandawate and Shah, 1969 | Tick | *H. intermedia* | India | Ganjam, Orissa | 19.387388 | 85.051544 | Goat |
| | Host | | India | Bhanjanagar, Orissa | 19.9358071 | 84.582481 | Goat |
| | Host | | India | Arehally, Mysore | 13.0462575 | 75.8040022 | Goat |
| | Host | | India | Zeban, Kashmir | 33.7125334 | 74.9866287 | Sheep |
| Joshi et al., 2005 | Tick | *H. intermedia*, *R. hemaphysaloides* | India | Manjari, Maharashtra | 19.4822619 | 74.7900264 | Goat |
| | Tick | *H. intermedia* | India | Saswad, Maharashtra | 18.3463128 | 74.0301826 | Sheep, Goat |
| | Tick | *H. intermedia* | India | Paud, Maharashtra | 18.5243308 | 73.6138821 | Goat |
| | Tick | *H. intermedia* | India | Loni, Maharashtra | 19.58041 | 74.4659065 | Goat |
| Boshell, 1970 | Tick | *H. intermedia* | India | Hubballi, Karnataka | 15.3647083 | 75.1239547 | Sheep, Goat |
| | Tick | *H. intermedia* | India | Barur, Tamil Nadu | 12.305112 | 78.305268 | Sheep, Goat |
| | Tick | *H. intermedia* | India | Sagar, Madhya Pradesh | 23.838805 | 78.7378068 | Sheep, Goat |
| Joshi et al., 1998 | Host | | India | Veerapuram, Tamil Nadu | 12.554789 | 80.093853 | Sheep, Goat, Human |
| Perera et al., 1996 | Tick, Host | *H. intermedia* | Sri Lanka | Kottukachchiya (Breeding Station) | 7.925625 | 79.969421 | Goat, Human |
| Weinbren et al., 1958 | Host | | Uganda | Kawanda | 0.422329 | 32.541517 | Sheep |
| | Host | | Uganda | Kamuli | 0.944785 | 33.126717 | Goat |
| | Host | | Uganda | Kampiringisa | 0.189173 | 32.246467 | Sheep |
| | Host | | Uganda | Mbarara | -0.60716 | 30.654502 | Sheep |
| | Host | | Uganda | Entebbe | 0.051184 | 32.463708 | Goat |
| Jesset, 1978 | Host | | Tanzania | Arusha | -3.386925 | 36.682993 | Sheep, Goat |
| | Host | | Tanzania | Mwanza | -2.51643 | 32.917452 | Sheep, Goat |
| | Host | | Tanzania | Mpwapwa | -6.347795 | 36.48512 | Sheep, Goat |
| | Host | | Tanzania | Dar-es-Salaam | -6.792354 | 39.208328 | Sheep, Goat |

| | | | | | | |
|---|---|---|---|---|---|---|
| | Host | Tanzania | Iringa | -7.773094 | 35.69912 | Sheep, Goat |
| | Host | Tanzania | Loliondo | -2.05 | 35.616667 | Sheep, Goat |
| | Host | Tanzania | Tabora | -5.042495 | 32.819733 | Sheep, Goat |
| Terpestra, 1969 | Host | Uganda | Okoro, West Nile | 2.5544293 | 30.9417368 | Sheep |
| | Host | Uganda | Ayivu, West Nile | 3.037925 | 30.9417368 | Goat |
| | Host | Uganda | Maracha, West Nile | 3.2873127 | 30.9403023 | Goat |
| | Host | Uganda | Terego, West Nile | 3.1710085 | 31.1252135 | Goat |
| | Host | Uganda | Koboko, West Nile | 3.4167992 | 30.9589496 | Sheep, Goat |
| | Host | Uganda | Aringa, West Nile | 3.4448673 | 31.1975236 | Goat |
| | Host | Uganda | Bujenje, Bunyoro | 1.5939148 | 31.5826642 | Sheep, Goat |
| | Host | Uganda | Buruli, Bunyoro | 1.3489721 | 32.4467238 | Sheep, Goat |
| | Host | Uganda | Bugahya, Bunyoro | 1.6343426 | 31.1710389 | Sheep, Goat |
| | Host | Uganda | Mwenge, Toro | 0.7137652 | 30.6199895 | Sheep, Goat |
| | Host | Uganda | Burahya, Toro | 0.6832535 | 30.2974199 | Sheep, Goat |
| | Host | Uganda | Bunyangabu, Toro | 0.4870918 | 30.2051096 | Sheep, Goat |
| | Host | Uganda | Busongora, Toro | 0.3244003 | 30.0665236 | Goat |
| | Host | Uganda | Bujumbura, Kigezi | -0.5546336 | 29.8352303 | Sheep, Goat |
| | Host | Uganda | Kinkizi, Kigezi | -0.8195253 | 29.742604 | Goat |
| | Host | Uganda | Rukiga, Kigezi | -1.1326337 | 30.043412 | Sheep, Goat |
| | Host | Uganda | Bufumbira, Kigezi | -1.1538569 | 29.6499162 | Sheep, Goat |
| | Host | Uganda | Ndorwa, Kigezi | -1.3504705 | 30.0202964 | Sheep, Goat |
| | Host | Uganda | Tororo, Bukedi | 0.6782274 | 34.1865669 | Sheep |
| | Host | Uganda | Samia Bugwe, Bukedi | 0.4044731 | 34.0195827 | Goat |
| Edelsten, 1975 | Host | Somalia | | 10.34 | 43.113 | Sheep, Goat |
| | Host | Somalia | | 10.286 | 43.443 | Sheep, Goat |
| | Host | Somalia | | 10.064 | 43.209 | Sheep, Goat |
| | Host | Somalia | | 9.907 | 43.513 | Sheep, Goat |
| | Host | Somalia | | 9.783 | 43.207 | Sheep, Goat |
| | Host | Somalia | | 9.741 | 43.773 | Sheep, Goat |
| | Host | Somalia | | 9.867 | 44.304 | Sheep, Goat |
| | Host | Somalia | | 9.476 | 43.504 | Sheep, Goat |
| | Host | Somalia | | 9.439 | 43.797 | Sheep, Goat |
| | Host | Somalia | | 9.203 | 43.935 | Sheep, Goat |
| | Host | Somalia | | 9.224 | 44.41 | Sheep, Goat |
| | Host | Somalia | | 8.969 | 44.391 | Sheep, Goat |
| | Host | Somalia | | 9.427 | 44.887 | Sheep, Goat |
| | Host | Somalia | | 9.9 | 44.782 | Sheep, Goat |
| | Host | Somalia | | 9.75 | 45.179 | Sheep, Goat |
| | Host | Somalia | | 8.833 | 44.824 | Sheep, Goat |
| | Host | Somalia | | 8.709 | 45.154 | Sheep, Goat |
| | Host | Somalia | | 9.481 | 45.348 | Sheep, Goat |
| | Host | Somalia | | 9.161 | 45.409 | Sheep, Goat |
| | Host | Somalia | | 8.489 | 45.776 | Sheep, Goat |

| | | | | | |
|---|---|---|---|---|---|
| | Host | Somalia | 8.667 | 46.01 | Sheep, Goat |
| | Host | Somalia | 9.041 | 46.347 | Sheep, Goat |
| | Host | Somalia | 8.358 | 46.143 | Sheep, Goat |
| | Host | Somalia | 8.693 | 46.513 | Sheep, Goat |
| | Host | Somalia | 9.259 | 46.679 | Sheep, Goat |
| | Host | Somalia | 9.554 | 47.126 | Sheep, Goat |
| | Host | Somalia | 8.744 | 47.051 | Sheep, Goat |
| | Host | Somalia | 8.281 | 46.679 | Sheep, Goat |
| | Host | Somalia | 8.576 | 47.484 | Sheep, Goat |
| | Host | Somalia | 8.077 | 47.119 | Sheep, Goat |
| | Host | Somalia | 8.07 | 47.875 | Sheep, Goat |
| | Host | Somalia | 8.487 | 48.085 | Sheep, Goat |
| | Host | Somalia | 9.238 | 48.34 | Sheep, Goat |
| | Host | Somalia | 9.862 | 48.614 | Sheep, Goat |
| | Host | Somalia | 9.659 | 49.115 | Sheep, Goat |
| Davies, 1978[1] | Host | Kenya | 1.494 | 35.524 | Sheep, Goat |
| | Host | Kenya | 1.384 | 35.517 | Sheep, Goat |
| | Host | Kenya | 1.346 | 35.402 | Sheep, Goat |
| | Host | Kenya | 1.247 | 35.27 | Sheep, Goat |
| | Host | Kenya | 1.315 | 35.616 | Sheep, Goat |
| | Host | Kenya | 1.205 | 35.699 | Sheep, Goat |
| | Host | Kenya | 1.111 | 35.69 | Sheep, Goat |
| | Host | Kenya | 0.885 | 35.569 | Sheep, Goat |
| | Host | Kenya | 0.788 | 35.562 | Sheep, Goat |
| | Host | Kenya | 0.758 | 35.43 | Sheep, Goat |
| | Host | Kenya | 0.676 | 35.572 | Sheep, Goat |
| | Host | Kenya | 0.581 | 35.574 | Sheep, Goat |
| | Host | Kenya | 0.452 | 35.6 | Sheep, Goat |
| | Host | Kenya | 0.376 | 35.442 | Sheep, Goat |
| | Host | Kenya | 0.771 | 35.314 | Sheep, Goat |
| | Host | Kenya | 0.606 | 35.257 | Sheep, Goat |
| | Host | Kenya | 0.685 | 34.573 | Sheep, Goat |
| | Host | Kenya | 0.716 | 34.755 | Sheep, Goat |
| | Host | Kenya | 0.635 | 34.803 | Sheep, Goat |
| | Host | Kenya | 0.043 | 34.341 | Sheep, Goat |
| | Host | Kenya | 0.165 | 34.556 | Sheep, Goat |
| | Host | Kenya | 0.19 | 35.009 | Sheep, Goat |
| | Host | Kenya | 0.399 | 35.19 | Sheep, Goat |
| | Host | Kenya | 0.336 | 35.165 | Sheep, Goat |
| | Host | Kenya | 0.264 | 35.156 | Sheep, Goat |
| | Host | Kenya | 0.218 | 35.193 | Sheep, Goat |

| | | | | |
|---|---|---|---|---|
| Host | Kenya | 0.138 | 35.152 | Sheep, Goat |
| Host | Kenya | 0.085 | 35.214 | Sheep, Goat |
| Host | Kenya | 0.173 | 35.285 | Sheep, Goat |
| Host | Kenya | 0.426 | 35.782 | Sheep, Goat |
| Host | Kenya | 0.276 | 35.839 | Sheep, Goat |
| Host | Kenya | -0.838 | 34.24 | Sheep, Goat |
| Host | Kenya | -0.718 | 34.286 | Sheep, Goat |
| Host | Kenya | -0.236 | 35.546 | Sheep, Goat |
| Host | Kenya | -0.235 | 35.665 | Sheep, Goat |
| Host | Kenya | -0.33 | 35.536 | Sheep, Goat |
| Host | Kenya | -0.506 | 34.859 | Sheep, Goat |
| Host | Kenya | -0.465 | 35.037 | Sheep, Goat |
| Host | Kenya | -0.494 | 35.192 | Sheep, Goat |
| Host | Kenya | -0.875 | 34.763 | Sheep, Goat |
| Host | Kenya | -0.807 | 34.831 | Sheep, Goat |
| Host | Kenya | -0.755 | 35.007 | Sheep, Goat |
| Host | Kenya | -0.826 | 35.138 | Sheep, Goat |
| Host | Kenya | -0.935 | 35.149 | Sheep, Goat |
| Host | Kenya | -0.948 | 35.016 | Sheep, Goat |
| Host | Kenya | -1.037 | 35.066 | Sheep, Goat |
| Host | Kenya | -1.127 | 34.873 | Sheep, Goat |
| Host | Kenya | -1.299 | 34.868 | Sheep, Goat |
| Host | Kenya | -1.107 | 35.101 | Sheep, Goat |
| Host | Kenya | -1.203 | 35.056 | Sheep, Goat |
| Host | Kenya | -1.287 | 35.185 | Sheep, Goat |
| Host | Kenya | -1.132 | 35.307 | Sheep, Goat |
| Host | Kenya | -1.312 | 35.321 | Sheep, Goat |
| Host | Kenya | -1.175 | 35.472 | Sheep, Goat |
| Host | Kenya | -0.966 | 35.501 | Sheep, Goat |
| Host | Kenya | -1 | 35.82 | Sheep, Goat |
| Host | Kenya | -1.049 | 36.039 | Sheep, Goat |
| Host | Kenya | -1.202 | 35.838 | Sheep, Goat |
| Host | Kenya | -1.544 | 35.639 | Sheep, Goat |
| Host | Kenya | -1.437 | 35.88 | Sheep, Goat |
| Host | Kenya | -1.827 | 35.727 | Sheep, Goat |
| Host | Kenya | -1.781 | 35.873 | Sheep, Goat |
| Host | Kenya | -0.114 | 36.82 | Sheep, Goat |
| Host | Kenya | -0.202 | 36.921 | Sheep, Goat |
| Host | Kenya | -0.399 | 36.918 | Sheep, Goat |
| Host | Kenya | -0.477 | 36.957 | Sheep, Goat |
| Host | Kenya | -0.56 | 36.918 | Sheep, Goat |

| | | | | |
|---|---|---|---|---|
| Host | Kenya | -0.738 | 36.5 | Sheep, Goat |
| Host | Kenya | -0.912 | 36.476 | Sheep, Goat |
| Host | Kenya | -0.902 | 36.695 | Sheep, Goat |
| Host | Kenya | -1.009 | 36.677 | Sheep, Goat |
| Host | Kenya | -0.967 | 36.774 | Sheep, Goat |
| Host | Kenya | -1.042 | 36.843 | Sheep, Goat |
| Host | Kenya | -1.084 | 36.789 | Sheep, Goat |
| Host | Kenya | -1.125 | 36.845 | Sheep, Goat |
| Host | Kenya | -1.184 | 36.877 | Sheep, Goat |
| Host | Kenya | -1.195 | 36.699 | Sheep, Goat |
| Host | Kenya | -1.048 | 37.01 | Sheep, Goat |
| Host | Kenya | -0.851 | 36.953 | Sheep, Goat |
| Host | Kenya | -0.96 | 37.064 | Sheep, Goat |
| Host | Kenya | -0.863 | 37.091 | Sheep, Goat |
| Host | Kenya | -0.7 | 36.954 | Sheep, Goat |
| Host | Kenya | -0.622 | 37.037 | Sheep, Goat |
| Host | Kenya | -0.725 | 37.077 | Sheep, Goat |
| Host | Kenya | -0.651 | 37.15 | Sheep, Goat |
| Host | Kenya | -0.559 | 37.184 | Sheep, Goat |
| Host | Kenya | -0.765 | 37.24 | Sheep, Goat |
| Host | Kenya | -0.559 | 37.26 | Sheep, Goat |
| Host | Kenya | -1.116 | 37.187 | Sheep, Goat |
| Host | Kenya | -1.174 | 37.087 | Sheep, Goat |
| Host | Kenya | -1.236 | 36.947 | Sheep, Goat |
| Host | Kenya | -1.228 | 37.044 | Sheep, Goat |
| Host | Kenya | -1.28 | 37.008 | Sheep, Goat |
| Host | Kenya | -1.394 | 36.728 | Sheep, Goat |
| Host | Kenya | -1.157 | 37.343 | Sheep, Goat |
| Host | Kenya | -1.518 | 37.051 | Sheep, Goat |
| Host | Kenya | -1.474 | 37.146 | Sheep, Goat |
| Host | Kenya | -1.498 | 37.346 | Sheep, Goat |
| Host | Kenya | -1.915 | 37.236 | Sheep, Goat |
| Host | Kenya | -0.809 | 37.327 | Sheep, Goat |
| Host | Kenya | -0.769 | 37.409 | Sheep, Goat |
| Host | Kenya | -0.657 | 37.376 | Sheep, Goat |
| Host | Kenya | -0.557 | 37.334 | Sheep, Goat |
| Host | Kenya | -0.582 | 37.51 | Sheep, Goat |
| Host | Kenya | -0.671 | 37.664 | Sheep, Goat |
| Host | Kenya | -0.759 | 37.712 | Sheep, Goat |
| Host | Kenya | -0.53 | 37.842 | Sheep, Goat |
| Host | Kenya | -0.482 | 37.634 | Sheep, Goat |

| | | | | |
|---|---|---|---|---|
| Host | Kenya | -0.387 | 37.714 | Sheep, Goat |
| Host | Kenya | -0.347 | 37.528 | Sheep, Goat |
| Host | Kenya | -0.299 | 37.691 | Sheep, Goat |
| Host | Kenya | -0.23 | 37.722 | Sheep, Goat |
| Host | Kenya | -0.183 | 37.638 | Sheep, Goat |
| Host | Kenya | -0.139 | 37.725 | Sheep, Goat |
| Host | Kenya | -0.072 | 37.693 | Sheep, Goat |
| Host | Kenya | -0.038 | 37.625 | Sheep, Goat |
| Host | Kenya | -0.004 | 37.696 | Sheep, Goat |
| Host | Kenya | -0.021 | 37.796 | Sheep, Goat |
| Host | Kenya | 0.004 | 37.864 | Sheep, Goat |
| Host | Kenya | -0.323 | 37.958 | Sheep, Goat |
| Host | Kenya | -0.387 | 38.062 | Sheep, Goat |
| Host | Kenya | 0.099 | 37.926 | Sheep, Goat |
| Host | Kenya | 0.127 | 37.825 | Sheep, Goat |
| Host | Kenya | 0.192 | 37.885 | Sheep, Goat |
| Host | Kenya | 0.192 | 37.993 | Sheep, Goat |
| Host | Kenya | 0.283 | 37.987 | Sheep, Goat |
| Host | Kenya | -2.927 | 37.486 | Sheep, Goat |
| Host | Kenya | -2.919 | 37.623 | Sheep, Goat |
| Host | Kenya | -3.052 | 37.715 | Sheep, Goat |
| Host | Kenya | -3.431 | 37.748 | Sheep, Goat |
| Host | Kenya | -3.395 | 37.903 | Sheep, Goat |
| Host | Kenya | -3.486 | 37.797 | Sheep, Goat |
| Host | Kenya | -3.466 | 38.049 | Sheep, Goat |
| Host | Kenya | -3.491 | 38.145 | Sheep, Goat |
| Host | Kenya | -3.55 | 38.333 | Sheep, Goat |
| Host | Kenya | -3.588 | 38.38 | Sheep, Goat |
| Host | Kenya | -3.62 | 38.461 | Sheep, Goat |
| Host | Kenya | -3.459 | 38.385 | Sheep, Goat |
| Host | Kenya | -3.402 | 38.455 | Sheep, Goat |
| Host | Kenya | -3.546 | 38.479 | Sheep, Goat |
| Host | Kenya | -3.566 | 38.602 | Sheep, Goat |
| Host | Kenya | -3.649 | 38.651 | Sheep, Goat |
| Host | Kenya | -3.819 | 38.7 | Sheep, Goat |
| Host | Kenya | -3.87 | 38.674 | Sheep, Goat |
| Host | Kenya | -3.886 | 38.754 | Sheep, Goat |
| Host | Kenya | -4.235 | 39.581 | Sheep, Goat |
| Host | Kenya | -4.01 | 39.616 | Sheep, Goat |
| Host | Kenya | -4.015 | 39.676 | Sheep, Goat |
| Host | Kenya | -3.961 | 39.701 | Sheep, Goat |

| | | | | | |
|---|---|---|---|---|---|
| | Host | Kenya | -3.891 | 39.735 | Sheep, Goat |
| | Host | Kenya | -3.759 | 39.79 | Sheep, Goat |
| | Host | Kenya | -3.646 | 39.561 | Sheep, Goat |
| | Host | Kenya | -3.63 | 39.777 | Sheep, Goat |
| | Host | Kenya | -3.588 | 39.852 | Sheep, Goat |
| | Host | Kenya | -3.458 | 39.905 | Sheep, Goat |
| Davies, 1978[2] | Host | Kenya | 0.282 | 35.034 | Sheep, Goat |
| | Host | Kenya | 0.163 | 35.097 | Sheep, Goat |
| | Host | Kenya | 0.617 | 35.097 | Sheep, Goat |
| | Host | Kenya | 0.536 | 35.253 | Sheep, Goat |
| | Host | Kenya | -0.096 | 36.74 | Sheep, Goat |
| | Host | Kenya | -0.187 | 36.957 | Sheep, Goat |
| | Host | Kenya | -0.288 | 36.698 | Sheep, Goat |
| | Host | Kenya | -0.428 | 37.164 | Sheep, Goat |
| | Host | Kenya | -1.147 | 36.355 | Sheep, Goat |
| | Host | Kenya | -0.929 | 36.513 | Sheep, Goat |
| | Host | Kenya | -0.879 | 36.617 | Sheep, Goat |
| | Host | Kenya | -1.249 | 36.515 | Sheep, Goat |
| | Host | Kenya | -1.36 | 36.55 | Sheep, Goat |
| | Host | Kenya | -1.293 | 36.607 | Sheep, Goat |
| | Host | Kenya | -1.191 | 36.619 | Sheep, Goat |
| | Host | Kenya | -1.086 | 36.601 | Sheep, Goat |
| | Host | Kenya | -1.13 | 36.713 | Sheep, Goat |
| | Host | Kenya | -1.239 | 36.7 | Sheep, Goat |
| | Host | Kenya | -1.345 | 36.693 | Sheep, Goat |
| | Host | Kenya | -1.027 | 36.739 | Sheep, Goat |
| | Host | Kenya | -0.977 | 36.829 | Sheep, Goat |
| | Host | Kenya | -1.088 | 36.817 | Sheep, Goat |
| | Host | Kenya | -1.205 | 36.798 | Sheep, Goat |
| | Host | Kenya | -1.301 | 36.777 | Sheep, Goat |
| | Host | Kenya | -1.161 | 36.881 | Sheep, Goat |
| | Host | Kenya | -1.059 | 36.916 | Sheep, Goat |
| | Host | Kenya | -0.844 | 36.998 | Sheep, Goat |
| | Host | Kenya | -0.979 | 36.996 | Sheep, Goat |
| | Host | Kenya | -1.11 | 36.993 | Sheep, Goat |
| | Host | Kenya | -1.046 | 37.095 | Sheep, Goat |
| | Host | Kenya | -1.191 | 37.084 | Sheep, Goat |
| | Host | Kenya | -1.206 | 36.957 | Sheep, Goat |
| | Host | Kenya | -1.28 | 36.88 | Sheep, Goat |
| | Host | Kenya | -1.379 | 36.836 | Sheep, Goat |
| | Host | Kenya | -1.42 | 36.772 | Sheep, Goat |

| | | | | |
|---|---|---|---|---|
| Host | Kenya | -1.359 | 36.938 | Sheep, Goat |
| Host | Kenya | -1.452 | 36.911 | Sheep, Goat |
| Host | Kenya | -1.442 | 36.992 | Sheep, Goat |
| Host | Kenya | -1.402 | 37.09 | Sheep, Goat |
| Host | Kenya | -1.312 | 37.134 | Sheep, Goat |
| Host | Kenya | -1.291 | 37.034 | Sheep, Goat |
| Host | Kenya | -0.969 | 37.24 | Sheep, Goat |
| Host | Kenya | -0.984 | 37.42 | Sheep, Goat |
| Host | Kenya | -1.647 | 37.185 | Sheep, Goat |
| Host | Kenya | -1.759 | 37.176 | Sheep, Goat |
| Host | Kenya | -1.775 | 37.297 | Sheep, Goat |
| Host | Kenya | -2.857 | 37.439 | Sheep, Goat |
| Host | Kenya | -2.886 | 37.578 | Sheep, Goat |
| Host | Kenya | -2.977 | 37.668 | Sheep, Goat |
| Host | Kenya | -4.271 | 39.529 | Sheep, Goat |
| Host | Kenya | -4.087 | 39.601 | Sheep, Goat |
| Host | Kenya | -4.034 | 39.483 | Sheep, Goat |
| Host | Kenya | -4.008 | 39.58 | Sheep, Goat |
| Host | Kenya | -3.988 | 39.673 | Sheep, Goat |
| Host | Kenya | -3.912 | 39.624 | Sheep, Goat |
| Host | Kenya | -3.915 | 39.727 | Sheep, Goat |
| Host | Kenya | -3.903 | 39.512 | Sheep, Goat |
| Host | Kenya | -3.873 | 39.411 | Sheep, Goat |
| Host | Kenya | -3.812 | 39.583 | Sheep, Goat |
| Host | Kenya | -3.827 | 39.671 | Sheep, Goat |
| Host | Kenya | -3.816 | 39.765 | Sheep, Goat |
| Host | Kenya | -3.7 | 39.573 | Sheep, Goat |
| Host | Kenya | -3.719 | 39.657 | Sheep, Goat |
| Host | Kenya | -3.78 | 39.474 | Sheep, Goat |
| Host | Kenya | -3.707 | 39.764 | Sheep, Goat |
| Host | Kenya | -3.615 | 39.642 | Sheep, Goat |
| Host | Kenya | -3.551 | 39.73 | Sheep, Goat |
| Host | Kenya | -3.609 | 39.821 | Sheep, Goat |
| Host | Kenya | -3.452 | 39.757 | Sheep, Goat |
| Host | Kenya | -3.469 | 39.877 | Sheep, Goat |

[1] Davies, F. G. "A survey of Nairobi sheep disease antibody in sheep and goats, wild ruminants and rodents within Kenya." Epidemiology & Infection 81.2 (1978): 251-258.

[2] Davies, F. G. "Nairobi sheep disease in Kenya. The isolation of virus from sheep and goats, ticks and possible maintenance hosts." Epidemiology &

Infection 81.2 (1978): 259-265.

**Table S2:** Systematic review process steps, as described by O’Connor and Sargeant, 2014.

| |
|---|
| **Pre-Step: Assemble review team and develop a systematic review protocol**<br>SK will be the main person responsible for compiling and reviewing all of the literature on NSDv. A review protocol will be prepared in accordance with O’Connor and Sargeant’s methods. |
| **Step 1: Define the review question**<br>For reviews about disease burden (prevalence or incidence), the format uses the acronym PO: P=Population and O=Outcome (E.F.S.A., 2010).<br><br>Our research question is as follows:<br>*What is the prevalence of Nairobi Sheep Disease in small ruminants, and is it possible to identify and anticipate vulnerable areas and populations?* |
| **Step 2: Conduct extensive search for studies**<br>The aim of conducting an extensive search is to ensure as many relevant results as possible are included in the review. The rationale is to reduce the bias associated with the accessibility of studies based on their outcome, sometimes called retrieval bias (O’Connor and Sargeant, 2014). In animal health and welfare reviews, the inclusion of CAB Abstracts is likely to be important, as data suggest that it provides the most comprehensive coverable of animal health topics (Grindlay et al., 2012). We chose to use EBSCOhost to search for our results because it includes CAB Abstracts.<br><br>***EBSCOhost (339 total results)***<br>*CAB Abstracts (125)*<br>*CAB Abstracts Archive (83)*<br>*MEDLINE (74)*<br>*Agricola (32)*<br>*Academic Search Complete (15)*<br>*CINAHL Plus with Full Text (2)*<br>*Environment Complete (2)*<br>*OpenDissertations (2)*<br>*Wildlife and Ecology Studies Worldwide (1)*<br>*MasterFILE Premier (1)*<br>*Military & Government Collection (1)* |
| **Step 3: Select relevant studies from the results of the search**<br>Once the citations have been retrieved, it is necessary to evaluate them and identify those relevant to the review. A series of short questions are applied to the title and abstract of each citation identified (O’Connor and Sargeant, 2014). Due to the small number of publications on NSDv, we wanted to review every publication on NSDv available. For this reason, we left our questions short and simple:<br>*Does the title or abstract mention Nairobi Sheep Disease or Ganjam Virus?* |
| **Step 4: Collecting data from relevant studies** |

| |
|---|
| In this step, the results of the relevant studies are extracted (O’Connor and Sargeant, 2014). We are interested in capturing disease distribution in order to build upon our ecological model, so we collected geographic locations of NSDv occurrences. |
| **Step 5: Assess the risk of bias in relevant studies**<br>It is important to report the potential for bias to the end user in order to alert them to potential concerns and uncertainties about the evidence summary. Since we will be including different study designs in our review, this information will be reported to the reader (O’Connor and Sargeant, 2014). |
| **Step 6: Synthesize the results**<br>Frequently in animal health, animal welfare, and food safety, it is not possible to combine the results in a meta-analysis. This often occurs due to a low number of relevant studies or because of poor reporting or differences in metrics used to measure the outcomes. Therefore, the presentation of results and discussion takes the place of this step (O’Connor and Sargeant, 2014). |
| **Step 7: Present the results**<br>The presentation of the results of the review should include the following components:<br>- The results of the search and study selection;<br>- Summary information about the characteristics of the studies identified as relevant to the review (including those that could not be included in a meta-analysis);<br>- Risk of bias assessment for the individual studies;<br>- Outcomes reported by the relevant studies;<br>- Risk of bias across the studies.<br><br>The PRISMA Statement should be strictly adhered to, and there should be a way to present a summary of the overall assessment of the body of work, such as an evidence profile. An evidence profile provides a structured means of summarizing the risk of bias for each outcome included in the systematic review. The issues considered include inconsistency, the risk of bias in the studies, indirectness, imprecision, and other considerations (O’Connor and Sargeant, 2014). |
| **Step 8: Interpret the results and discussion**<br>Conclusions about the results of the review and a discussion of potential biases of the review should be discussed. PRISMA guidelines recommend the discussion and interpretation include a summary of the evidence, a discussion of the limitations of the review, and the overall conclusions (O’Connor and Sargeant, 2014). |

**Table S3:** PRISMA checklist (Moher et al., 2009).

| Section/topic | # | Checklist item | Reported on page # |
|---|---|---|---|
| **TITLE** | | | |
| Title | 1 | Identify the report as a systematic review, meta-analysis, or both. | |
| **ABSTRACT** | | | |
| Structured summary | 2 | Provide a structured summary including, as applicable: background; objectives; data sources; study eligibility criteria, participants, and interventions; study appraisal and synthesis methods; results; limitations; conclusions and implications of key findings; systematic review registration number. | 1,2 |
| **INTRODUCTION** | | | |
| Rationale | 3 | Describe the rationale for the review in the context of what is already known. | 8 |
| Objectives | 4 | Provide an explicit statement of questions being addressed with reference to participants, interventions, comparisons, outcomes, and study design (PICOS). | Table S2 |
| **METHODS** | | | |
| Protocol and registration | 5 | Indicate if a review protocol exists, if and where it can be accessed (e.g., Web address), and, if available, provide registration information including registration number. | NA |
| Eligibility criteria | 6 | Specify study characteristics (e.g., PICOS, length of follow-up) and report characteristics (e.g., years considered, language, publication status) used as criteria for eligibility, giving rationale. | Figure 2, Table S2 |
| Information sources | 7 | Describe all information sources (e.g., databases with dates of coverage, contact with study authors to identify additional studies) in the search and date last searched. | 9 |
| Search | 8 | Present full electronic search strategy for at least one database, including any limits used, such that it could be repeated. | 8 |
| Study selection | 9 | State the process for selecting studies (i.e., screening, eligibility, included in systematic review, and, if applicable, included in the meta-analysis). | 9 |
| Data collection process | 10 | Describe method of data extraction from reports (e.g., piloted forms, independently, in duplicate) and any processes for obtaining and confirming data from investigators. | 9 |
| Data items | 11 | List and define all variables for which data were sought (e.g., PICOS, funding sources) and any assumptions and simplifications made. | NA |
| Risk of bias in individual studies | 12 | Describe methods used for assessing risk of bias of individual studies (including specification of whether this was done at the study or outcome level), and how this information is to be used in any data synthesis. | NA |
| Summary measures | 13 | State the principal summary measures (e.g., risk ratio, difference in means). | NA |
| Synthesis of results | 14 | Describe the methods of handling data and combining results of studies, if done, including measures of consistency (e.g., $I^2$) for each meta-analysis. | NA |
| Risk of bias across studies | 15 | Specify any assessment of risk of bias that may affect the cumulative evidence (e.g., publication bias, selective reporting within studies). | NA |
| Additional analyses | 16 | Describe methods of additional analyses (e.g., sensitivity or subgroup analyses, meta-regression), if done, indicating which were pre-specified. | NA |
| **RESULTS** | | | |

| | | | |
|---|---|---|---|
| Study selection | 17 | Give numbers of studies screened, assessed for eligibility, and included in the review, with reasons for exclusions at each stage, ideally with a flow diagram. | Figure 2 |
| Study characteristics | 18 | For each study, present characteristics for which data were extracted (e.g., study size, PICOS, follow-up period) and provide the citations. | Table S1 |
| Risk of bias within studies | 19 | Present data on risk of bias of each study and, if available, any outcome level assessment (see item 12). | NA |
| Results of individual studies | 20 | For all outcomes considered (benefits or harms), present, for each study: (a) simple summary data for each intervention group (b) effect estimates and confidence intervals, ideally with a forest plot. | NA |
| Synthesis of results | 21 | Present the main results of the review. If meta-analyses are done, include for each, confidence intervals and measures of consistency" in accordance with the text in the Explanation and Elaboration document. | 13 |
| Risk of bias across studies | 22 | Present results of any assessment of risk of bias across studies (see Item 15). | NA |
| Additional analysis | 23 | Give results of additional analyses, if done (e.g., sensitivity or subgroup analyses, meta-regression [see Item 16]). | NA |
| **DISCUSSION** | | | |
| Summary of evidence | 24 | Summarize the main findings including the strength of evidence for each main outcome; consider their relevance to key groups (e.g., healthcare providers, users, and policy makers). | 17 |
| Limitations | 25 | Discuss limitations at study and outcome level (e.g., risk of bias), and at review-level (e.g., incomplete retrieval of identified research, reporting bias). | 19 |
| Conclusions | 26 | Provide a general interpretation of the results in the context of other evidence, and implications for future research. | 20 |
| **FUNDING** | | | |
| Funding | 27 | Describe sources of funding for the systematic review and other support (e.g., supply of data); role of funders for the systematic review. | 21 |
